# Liquid Phase Backscattered Scanning Electron Microscopy of *Bacillus subtilis* Spores

**DOI:** 10.64898/2026.03.24.713888

**Authors:** Jack Bromley, Adrián Pedrazo-Tardajos, Yongqiang Meng, Matthew C. Spink, Dogan Ozkaya, Rodney S. Ruoff, Graham Christie, Angus I. Kirkland, Judy S. Kim

## Abstract

Backscattered electron scanning electron microscopy (BSE-SEM) provides compositional image contrast but has found limited application to biological samples due to the low atomic number difference between constituent elements, the thickness of the surrounding environment, and the need for complex sample preparation. Here, we demonstrate the use of room temperature liquid phase BSE-SEM (LPBSEM) for imaging *Bacillus subtilis* spores encapsulated in graphene liquid cells, preserving native hydration and reducing the thickness of the sample environment. This approach eliminates the need for staining and enables high-contrast visualisation of subcellular structures. Distinct structural layers within *B. subtilis* spores have been observed with a contrast similar to conventional thin-section transmission electron microscopy but without the need for sample preparation that potentially compromises sample integrity. We further investigate the influence of beam energy on the interaction volume depth and image contrast and propose optimal conditions for subsurface visualisation. Monte Carlo simulations have been used to validate our experimental observations and provide a quantitative framework for understanding BSE generation from hydrated, low atomic number specimens.

## Introduction

Transmission electron microscopy (TEM)^1^ and scanning electron microscopy (SEM)^2^, have been used extensively to visualise biological structures at the molecular scale^3–6^. However, TEM requires thin sections and possibly heavy metal staining or focused ion beam (FIB) milling^7^, both of which risk introducing artifacts during sample preparation^7–10^. In contrast, SEM enables nanometre resolution surface-sensitive imaging through two main signals^2,10^. The secondary electron (SE) signal is sensitive to topographical features, whereas the backscattered electron (BSE) signal provides compositional contrast.

Despite this, BSE-SEM has not been frequently used for biological specimens^11^ due to the inherently low atomic number difference (Z) of the constituent light elements. To overcome this, contrast enhancement using immunogold labelling^12^ or osmium tetroxide staining^13^, have been used to increase local intensity of selectively stained features but can also alter or obscure ultrastructural detail^9^. Alternative preparation techniques such as FIB milling^14^ have been used but potentially introduce sample damage. As an example of the former, a recent study has demonstrated BSE-SEM imaging of neuronal cells with heavy metal staining in combination with correlative immunofluorescence techniques^15^.

A disadvantage of typical sample preparation for BSE-SEM of biological samples is that the signal originating from the surrounding medium (such as an embedding resin or vitreous ice in cryogenic preparations) significantly contributes to the overall image contrast^16,17^. Hence, due to the relatively high thickness of these media and concomitant background noise contributions the image signal-to-noise ratio (SNR) diminishes. This makes it difficult to resolve fine structural details within the specimen and can consequently obscure subtle variations in ultrastructure.

Advances in liquid-phase electron microscopy^18–23^ have provided new opportunities for imaging hydrated biological specimens without the need for heavy metal staining or dehydration using *in situ* liquid holders^24^, liquid chambers with silicon nitride windows^20,21,25^, and graphene liquid cells (GLCs)^18,19^. GLCs^22,26^ offer a particularly promising approach by encapsulating samples within a thin, electron-transparent membrane, effectively protecting and stabilising samples in a near native state while also minimising beam induced damage and charging artifacts^23,27,28^. Whilst liquid-phase TEM has been extensively explored^18,19^, and environmental SEM (ESEM) is well developed^29–31^ but requires a dedicated microscope configuration and typically involves resolution trade-offs^32^.

We report a sample preparation method that permits liquid phase SEM imaging in a standard instrument. In GLCs, the thin graphene membrane allows the electron beam to penetrate the sample with minimal scattering or absorption. The backscattered electrons generated can then escape through the thin layer graphene layer, preserving spatial resolution and improving the signal-to-noise ratio.

In this study, a workflow for liquid-phase BSE-SEM (LPBSEM) at room temperature in the SEM for imaging thick biological samples encapsulated within GLCs is described. As a case study, we use *Bacillus subtilis (B. subtilis)* spores^33,34^ which form when the bacteria are starved of nutrients and represent one of the most resilient biological structures found in nature, capable of withstanding extreme heat, desiccation, radiation, chemical stressors, and enzymatic degradation^35,36^. *B. subtilis* spores are also an ideal model species for this study due to their well characterised multilayered structure^37,38^. The overall structure of the *B. subtilis* spore has been extensively studied by negative stained thin section TEM, and it is now understood that the spore has a dense core surrounded by layers of concentric rings, namely the cortex (a thick layer of peptidoglycan) and the spore coat^39,40^. The spore coat can be further differentiated as two distinct proteinaceous sub layers; the inner coat and the outer coat. In some instances, the outermost, glycan-rich region of the outer coat, the crust, has been observed using TEM, but this has only been possible by sectioning and using Ruthenium Red^41^ as a glycan-specific stain which results in a loss of fine ultrastructural detail^42,43^. Native characterisation of spore structure is of significant interest in microbiology, as structural organisation is a key determinant for the environmental resistance of spores. There is also growing industrial interest driven by spore surface display technology, in which enzymes or other functional proteins are anchored to the spore surface for applications in biocatalysis, bioremediation, and antigen presentation^44–46^.

Each spore layer has a different composition. The coat is composed of more than 80 proteins^47^ and consists predominantly of C, H, O, and N, with smaller amounts of S and P from disulfide bonds and protein phosphorylation^48^, respectively. The cortex is a peptidoglycan-rich layer containing D-amino acids and muramic lactam^49^, while the core contains DNA, RNA, enzymes, and high levels of calcium dipicolinic acid (CaDPA)^37^. These compositional differences not only define the biological functions of each layer but also strongly affect how the layers interact with the incident electron beam, making the spore a suitably complex biological model for investigating imaging contrast in hydrated samples.

Using BSE imaging and the hydration-preserving properties of GLCs, we demonstrate the visualisation of distinct subcellular layers of the *B. subtilis* spore without staining, sectioning or FIB milling. We have also studied the effects of the incident beam energy on penetration depth and image contrast to optimise imaging parameters for subsurface visualisation. Monte Carlo simulations using CASINO V2.51^50^ provide insights into the BSE interaction volume within hydrated biological specimens as a validation of our experimental observations.

## Results & Discussion

### Effect of Hydration on Structural Integrity & Contrast

To compare hydrated and dehydrated spores, a spore-containing solution was dispensed on a graphene TEM grid (Figures 2D, E). The spores imaged using conventional SEM and thus dehydrated by the column vacuum, exhibited morphological collapse and loss of ultrastructural details due to dehydration and shrinkage. In contrast, encapsulating the *B. subtilis* spores in GLCs (Figure 2A, B) preserved the hydrated morphology, allowing visualisation of distinct subcellular layers.

**Figure 1.**
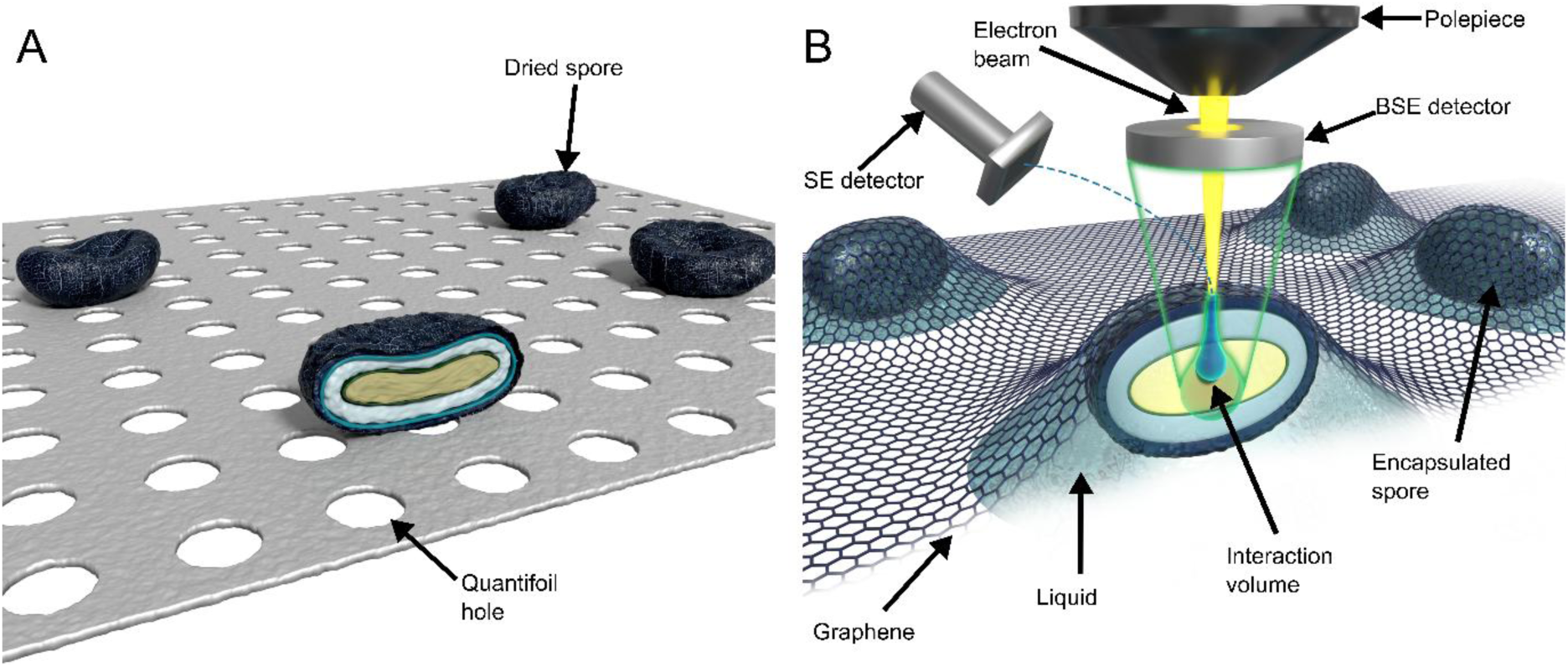
Schematic of a graphene-encapsulated *Bacillus subtilis* spore imaged using conventional room temperature scanning electron microscopy (A) and following graphene encapsulation (B) (the bottom graphene layer is not shown for clarity). Under dry conditions, the spore is exposed to the column vacuum, which is leads to dehydration and partial structural degradation or deformation of the internal architecture. When the spore is encapsulated between two graphene layers, it is surrounded by a thin layer of liquid sealed between these layers. The hexagonal lattice represents the top graphene membrane covering the spore. Due to the impermeability of graphene, the sample is not dehydrated as in A. B: Cross section of a spore with an illustration of the interaction volume of the incident electron beam. The larger green section represents the region where backscattered electrons are generated, and the smaller blue section represents the region where secondary electrons are generated.

**Figure 2:**
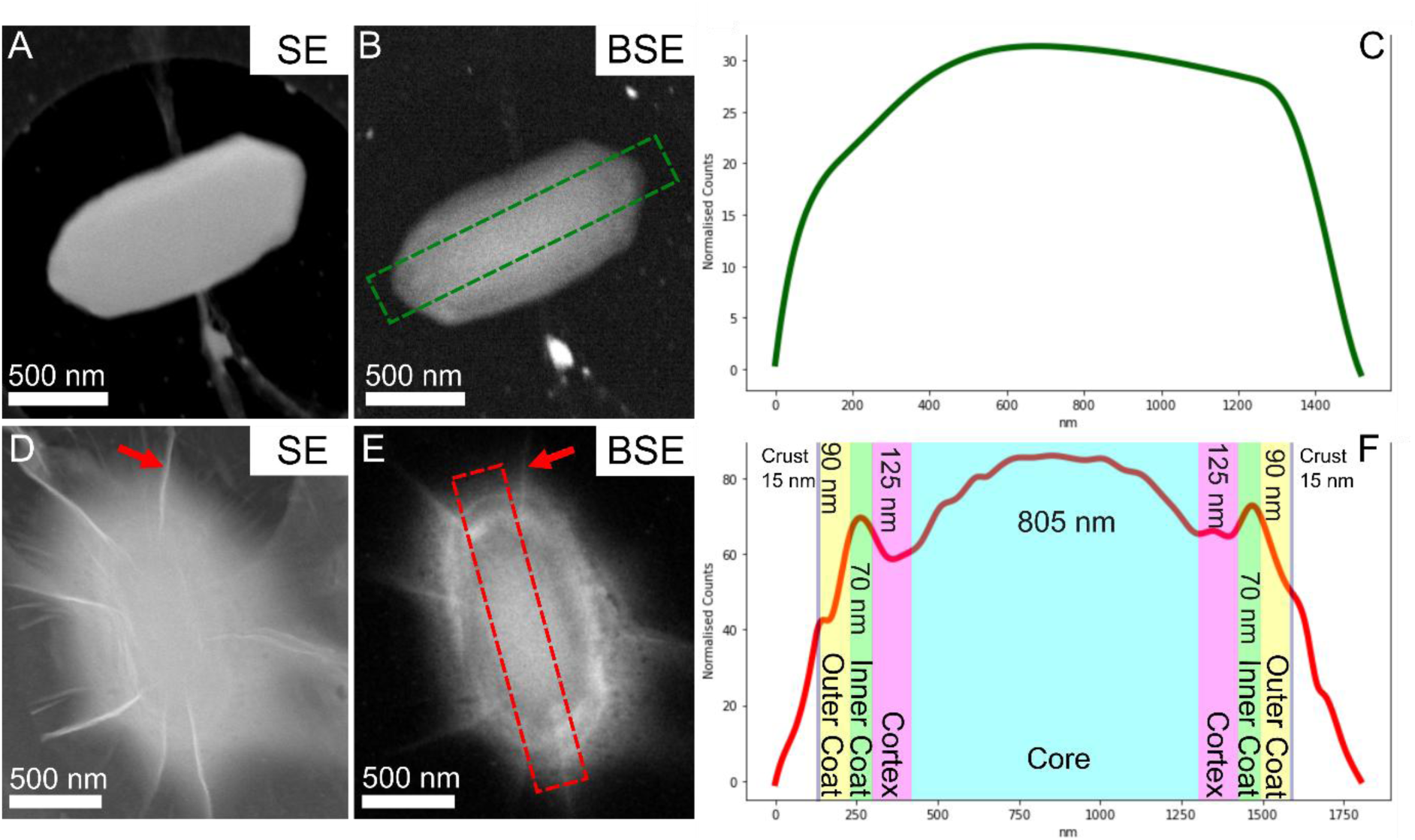
Comparison of SE and BSE imaging of *Bacillus subtilis (B. subtilis)* spores. (A-B) Dry and (D-E) wet environments imaged at 10 kV. D and E show a spore encapsulated with phosphate buffer (PB). Detailed internal features can only be seen in the BSE images when a hydrated environment is used. Line profiles of the BSE signal across the length of the respective spore (C) dry and (F) encapsulated in PB. Line profiles used a 40-pixel wide ROI. Red arrows in panels D and E indicate creases in the graphene.

Intensity profiles extracted across specific regions of interest in the BSE images (Figures 2C, F), show changes in intensity (and hence backscatter yield) which could be correlated with either the local density or elemental composition from region to region in the spore. Since the *B. subtilis* spore is an organic, biological material, there is limited variation in elemental composition and therefore the intensity and contrast variations are likely due to changes in density. Line profiles taken from the images allow direct comparison between the hydrated and dehydrated spores, illustrating how the use of GLCs maintain and preserve the structural integrity of the spore (Figure 2). Dehydrated spores displayed uniform low contrast, with no internal structural differentiation whereas, spores encapsulated in PB buffer showed contrast variations, with distinct peaks corresponding to the core, cortex, and spore coat. Measurement of the approximate thicknesses of *B. subtilis* spore layers gives values for: crust ∼15 nm, outer coat ∼90 nm, inner coat ∼70 nm, cortex ∼125 nm, and core diameter ∼805 nm along the long axis. The crust has previously only been observed in TEM studies of stained sample sections (with Ruthenium Red)^41^. Here, a distinct bright, membrane-like structure ∼15 nm in thickness is visible (Figure 2E). Given that the adjacent outer coat has a lower intensity, we interpret this bright layer as the crust. This reflects hydrated-state dimensions, which can be underestimated in conventional dehydrated/stained EM preparations^51^ since the cortex and coat are hydrated in the native state. It has been shown that the outer coat thickness has dimensions from 70 to 200 nm^52^ and along the short axis, our measurements are consistent with previous reports^53^ with a total coat thickness of ∼100 nm, cortex ∼90 nm thick, and core ∼475 nm thick. Along the short axis, the crust could not be directly visualised as the local density of the inner and outer coat in this region is higher, making it more difficult to differentiate these features. This indicates that at the poles of the spore, the coat is less tightly packed, whereas the cortex has a constant thickness and packing density. In turn this suggests that the hydration state affects the backscatter yield, as differences in local density become more pronounced when structural integrity is preserved.

Although electron-beam damage can occur in hydrated biological specimens at low fluences, its impact under the imaging conditions used here was evaluated by calculating the total electron fluence and by monitoring morphological changes during repeated imaging (Supplementary Information: Table S1 and Figure S1).

From line profile measurements, it has been determined that the length of the dry spore is 1305 ±52 nm (n = 10) and that of the PB encapsulated spore is 1486 nm ±57 (n = 10). Whilst the spore shown in Figure 2D and Figure 2E is encapsulated in phosphate buffer (PB), we have also encapsulated spores in pure water (Supplementary Information: Figure S2) to confirm that the observed contrast does not result from the buffer.

SE and BSE images (Figure 2D and 2E) of encapsulated spores were also compared to evaluate the effectiveness of BSE imaging in capturing subsurface detail. In whole spore samples, SE imaging revealed surface topology, with minimal differentiation between internal layers. In SE imaging the liquid layer between the spore and the graphene contributes to the SE signal, obscuring subsurface detail. In contrast, BSE imaging provided enhanced depth information, revealing subsurface features that are indiscernible in SE images as the BSEs have higher energy and are not absorbed substantially by the liquid layer. Importantly, the monoatomic graphene makes its contribution to the BSE signal minimal. Given the small interaction volume of the incident electrons with graphene, previous studies have reported an SE yield ratio of only ∼0.1 for graphene at 1 keV^54^. Since BSEs are known to originate deeper in the sample than SEs, the BSE yield is expected to be lower, (1.2x10^-4^ at 5 keV) (Supplementary Information: Table S2). However, there is a benefit to collecting both the SE and BSE signals simultaneously as although the liquid layer obscures spore detail in the SE signal, it can be used to confirm whether the sample is encapsulated since the coverage of the graphene can be directly visualised (Supplementary Information: Figure S3). The graphene coverage can further be assessed using the SE signal by taking low dose, low magnification images (1.8 e/Å^2^ and 8kX, respectively) to target areas of interest (Supplementary Information: Figure S4). It is important to note that charging was not observed during imaging of graphene encapsulated spores. Charging represents a significant barrier to SEM imaging in SE mode and whilst different strategies such as interleaved scanning can be used to reduce charging effects^55^, it has been reported that the high electrical conductivity of graphene reduces charging at low voltages^56^.

### Effect of Beam Energy on Penetration Depth & Image Contrast

The beam energy in LPBSEM influences the electron penetration depth and the interaction volume from which the BSE electrons are generated. Lower beam energies reduce electron penetration and the interaction volume, increasing surface sensitivity and resolution for a fixed sample tilt, whereas higher beam energies enable deeper penetration but reduce lateral resolution due to a larger interaction volume. Using the Kanaya-Okayama formula^57^ which accounts for both elastic scattering and inelastic energy losses the electron penetration depth estimated assuming a carbonaceous material with properties approximated by graphite (atomic number Z = 6, atomic weight A = 12.01 g mol⁻¹, and density ρ = 2.26 g cm⁻³) gives an estimated penetration depth of∼440 nm at 5 keV and ∼1400 nm at 10 keV.

For the Monte Carlo simulations, each layer of the spore was synthetically built based on thickness, composition, and density (See methods section). For simplicity, the spore coat was modelled as a single layer rather than subdivided into the inner coat, outer coat, and crust. This trend highlights how increasing beam energy shifts the origin of the BSE signal deeper into the sample, with lower energies offering better surface and edge definition, whereas higher energies probe internal composition. These findings further support the concept of a “quasi-cross section,” whereby contrast can be tuned using the beam energy to target specific layers of interest within a sample.

Figure 3 illustrates this at 5, 10, and 15 keV. At 5 keV, there is a clear distinction between the inner and outer coats, particularly at the spore poles. However, the core is dark and poorly resolved since BSEs are sparse in the core as confirmed by Monte Carlo simulations using CASINO^50^ (Figure 3(G)). At increased beam energy, the core is brighter, while the coat boundaries become less distinct. It is also worth noting that each spore within the GLCs is slightly different due to the graphene pockets containing a different amount of buffer/liquid.

**Figure 3:**
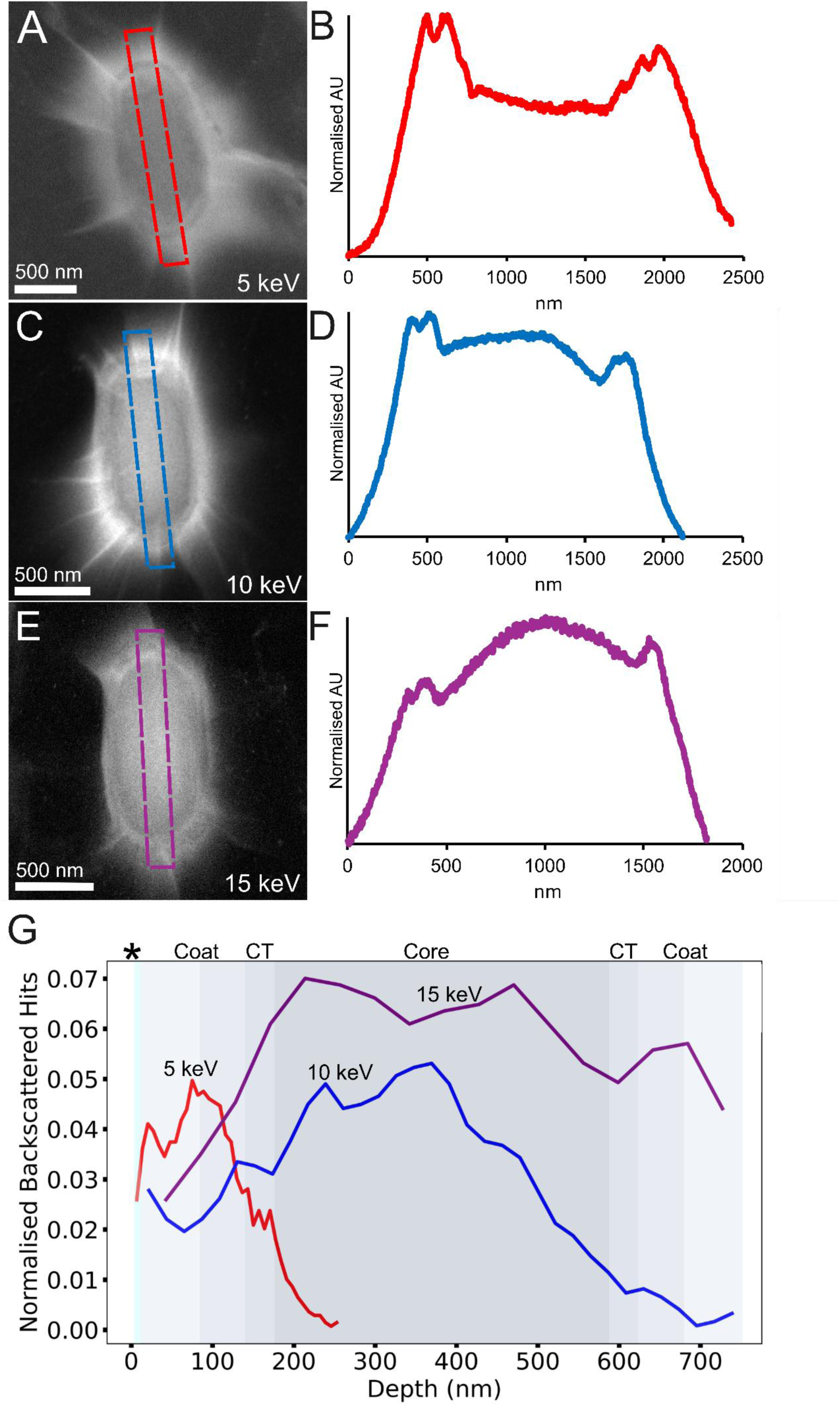
Liquid-phase backscattered electron SEM images of *Bacillus subtilis* spores. **(A) 5 keV (C) 10 keV, and (E) 15 keV.** Dashed rectangles indicate regions from which intensity line profiles were extracted, shown in **(B)** (red, 5 keV), **(D)** (blue, 10 keV), and **(F)** (purple, 15 keV). **(G)** Monte Carlo simulations of normalised backscattered electron generation as a function of depth. Shaded regions correspond to key spore structural layers, and curves indicate the relative contribution to signal by depth: red 5 keV, blue 10 keV, and purple 15 keV. Line profile ROI was 54 pixels for all beam energies. Increased beam energy leads to deeper penetration and broader signal distribution across spore layers. * = water, CT = cortex. The coat label spans two shades representing the inner and outer coat.

As already discussed, each layer of the spore has a different composition. In order to understand the effect the different layer compositions have on the image intensity, Monte Carlo simulations^50^ were used to model the BSE yield as a function of depth for different incident energies. For these calculations we considered the density and composition of each layer of the spore. There are no direct measurements of the density of the spore coat itself; however, helium displacement measurements of whole dry spores give a density of 1.33 g/cm^3^ ^58^, similar to that of dry proteinaceous material. Using this value gives estimates for the spore coat density of ∼1.3–1.4 g/cm³ for an average atomic number of ∼3.8 and atomic mass of ∼7.1amu (Table 2), values that are typical for proteinaceous material predominantly containing light elements^47^. As an example for CotA (an abundant protein in the spore coat^59^) as a representative protein, the elemental composition (∼54% C, 7% H, 17% N, 21% O, 1% S) by weight would yield an average atomic number of ∼3.8 and atomic mass ∼7.1. Note that post-translational modifications are not accounted for in this estimate.

**Table 1:** Measured lengths of the internal spore features along the long axis and total length in dry and encapsulated conditions.

| Feature | Size (nm) |
| --- | --- |
| Outer Coat | 90 |
| Inner Coat | 70 |
| Cortex | 125 |
| Core | 805 |
| Length (encapsulated, n =10) | 1486 |
| Length (dry, n =10) | 1305 |

**Table 2:** Estimated atomic weights, numbers and density for spore layers.

| Layer | Average atomic weight (g/mol) | Average Atomic Number (Z) | Density (ρ) (g/cm<sup>3</sup>) |
| --- | --- | --- | --- |
| Coat | ~7.1 | ~3.8 | 1.3 |
| Cortex | ~6.8 | ~3.7 | 1.2 |
| Core | ~7.4 | ~3.9 | 1.4 |

Inside the spore coat is the cortex, a peptidoglycan layer rich in D-amino acids and muramic lactam^49^. The cortex also contains low-Z elements with little P or S but more O^37^. There are no reported direct measurements of the cortex composition, although it is known to be a hydrated peptidoglycan with some protein crosslinking. Therefore, for our simulations we assumed a composition of 60% peptidoglycan, ∼30% water, and ∼10% protein. The elemental composition of peptidoglycan was estimated from reported analyses of bacterial cell wall material^60,61^. It consists of glycan chains of N-acetylglucosamine (GlcNAc; C₈H₁₅NO₆) and N-acetylmuramic acid (MurNAc; C₉H₁₇NO₇) crosslinked by short peptides rich in D-amino acids such as alanine and glutamic acid. A representative GlcNAc–MurNAc disaccharide with a four-to five–amino acid peptide stems yield an average composition of ∼44% carbon, 7% hydrogen, 8% nitrogen, and 41% oxygen by weight. With the additional assumed water and protein content included, this gives an estimated average atomic number of 3.7 and an atomic weight of 6.8 amu. Whilst polysaccharide density is generally higher than that of protein, the cortex is hydrated which reduces its density to approximately 1.2 g/cm³.

The core contains DNA, RNA, enzymes, and high levels of CaDPA, with CaDPA comprising ∼20% of the core mass (dry weight) and proteins ∼45% (dry). DNA contributes only a few percent of core mass, and water makes up ∼35% of the core wet weight. Based on these proportions, and using representative elemental compositions for CaDPA, protein, DNA, and water, the core is enriched in C, H, N, O, with significant contributions from Ca and P. This yields an estimated average atomic number of ∼4.0 and an average atomic mass of ∼7.5, reflecting the higher Ca and P content relative to the cortex and coat. Its density (∼1.4 g/cm³) is correspondingly higher, consistent with direct measurements showing that the core is much more dense than a typical bacterial protoplast^49^.

As the whole spore is analysed in the SEM, we have estimated the electron penetration depth for a better interpretation of our data. The electron penetration depth *R*, can be approximated using the Kanaya-Okayama formula^57^ (Eq. 1): 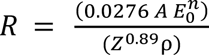 where *A* is the atomic weight in g/mole, *n* is a constant often chosen as ∼1.67, *E_0_* is the incident beam energy in kV, *Z* is the atomic number, and *ρ* is the density in g/cm^3^. This provides a useful analytical model for the interaction volume, demonstrating that higher accelerating voltages increase penetration depth, while higher atomic number and density reduce it. However, it should be noted that this model of electron penetration depth can become unreliable below 5 keV^62^. Assuming that the incident beam is not anomalously scattered at layer boundaries, we use an approximated average atomic weight of 12.5 g/mol, average atomic number of 7.5, and known density of 1.33 g/cm^3^ to estimate beam penetrations at various accelerating voltages in Table 3.

**Table 3:** Electron penetration depths (*R)* for the whole spore at various beam energies. *R* is calculated using the Kanaya-Okayama formula^57^.

| Accelerating Voltage ($E_0$ keV) | Electron Penetration Depth ($R$, nm) |
| --- | --- |
| 5 | 635 |
| 10 | 2019 |
| 15 | 3974 |

**Table 4:** Sample composition parameters used in CASINO for Monte Carlo Simulation. Thicknesses are based on short axis measurements in the results section. Compositions are based on representative proteins^59^ and inferences from previous data^39,49,60,61^.

| Sample Composition |  |  |  |  |  |  |  |  |
| --- | --- | --- | --- | --- | --- | --- | --- | --- |
| Layer & Thickness (nm) | C (% by weight) | H (% by weight) | O (% by weight) | N (% by weight) | S (% by weight) | P (% by weight) | Ca (% by weight) | Density (g/cm<sup>3</sup>) |
| Water (10 nm) | - | 11 | 89 | - | - | - | - | 1.0 |
| Coat (100 nm) | 54 | 7 | 17 | 21 | 1 | - | - | 1.3 |
| Cortex (90 nm) | 43 | 7 | 40 | 10 | - | - | - | 1.2 |
| Core (475 nm) | 34 | 7 | 43.9 | 12 | - | 0.1 | 3 | 1.4 |
| Cortex (90 nm) | 43 | 7 | 40 | 10 | - | - | - | 1.2 |
| Coat (100 nm) | 54 | 7 | 17 | 21 | 1 | - | - | 1.3 |

Whilst the Kanaya-Okayama formula is less accurate below 5 keV, it provides a useful estimate of penetration depth and allows for easy comparison between different beam energies. These suggest that since the full spore thickness is ∼ 750 - 1000 nm, beam energies above 5 keV interact with the entire volume, while lower energy electrons (< 5 keV) do not fully penetrate.

We note that penetration depth alone does not determine the image intensity. The penetration depth defines the interaction volume, but this is not specific to backscattered electron generation. Thus, we have investigated the depth at which backscattered electrons are generated using Monte Carlo simulations in CASINO^50^ (Figure 3G). At 5 kV a strong surface-weighted BSE signal is evident, peaking sharply in the outer and inner coats and diminishing rapidly past ∼200 nm. This is consistent with a limited penetration (around 635 nm) and strong surface sensitivity, emphasising outer features such as coat boundaries. At 10 kV the backscatter profile broadens, with the maximum signal arising from the cortex and the upper core region. At 15 kV backscatter generation is highest and includes signal from the core across the entire spore depth. However, despite the overall increase in BSE generation at 15 keV, the relative contribution from layers closer to the surface is reduced (Figure 3G).

### *Bacillus subtilis* Spore Germination –Snapshots into A Dynamic Process

Spore germination in *B. subtilis* follows a highly conserved and temporally ordered sequence of events that transition the spore from a metabolically dormant state to a germinated state with a loss of the characteristic, protective structure^63,64^. As previously reported, *B. subtilis* spores are deactivated under electron irradiation^65^, so germination was conducted *ex situ*. To capture structural intermediates that accompany these transitions, *B. subtilis* spores were germinated using a standard L-alanine-based protocol^66^ and imaged at various timepoints (based on optical microscope images Supplementary Information Figure S5) using LPBSEM. While graphene encapsulation does not permit real-time observation of single spores, it enables visualisation of internal changes in hydrated, unstained biological samples at different timepoints, preserving key features. Samples of germinating spores were encapsulated at various timepoints following germinant exposure, spanning the full range of the germination process (Supplementary Information Figure S5). Prior to encapsulation and imaging, each sample was screened using phase-contrast light microscopy to assess hallmark optical transitions, such as the shift from phase-bright (dormant) to phase-dark (germinated) spores, as a proxy for cortex degradation. Dormant spores appear phase-bright due to a dehydrated core and the presence of CaDPA, which give them a high refractive index relative to the surrounding medium. During germination, release of CaDPA and uptake of water into the spore core reduces the refractive index and the spores appears phase dark^67^. These early events also activate cortex-lytic enzymes (SleB and CwlJ), which degrade the peptidoglycan cortex and cause further core swelling and metabolic reactivation^49,68^. However, recent evidence has highlighted electron transport activity detected earlier than this point^69^. Ultimately, core swelling, increased turgor pressure, and rupture of the spore coat, allow the emergence and elongation of the vegetative cell^70^. In this study, alanine was used as the sole germinant and spores did not reach outgrowth but were arrested at the germinated spore stage.

In their dormant state, spores appear as highly ordered, multilayered structures (Figure 4 (1)) with distinct contrast differences between the spore coat, cortex, and core, visible in the BSE images (Figure 4(1), and Figure 2E which are the same spore). This marks the starting point for the germination process.

**Figure 4:**
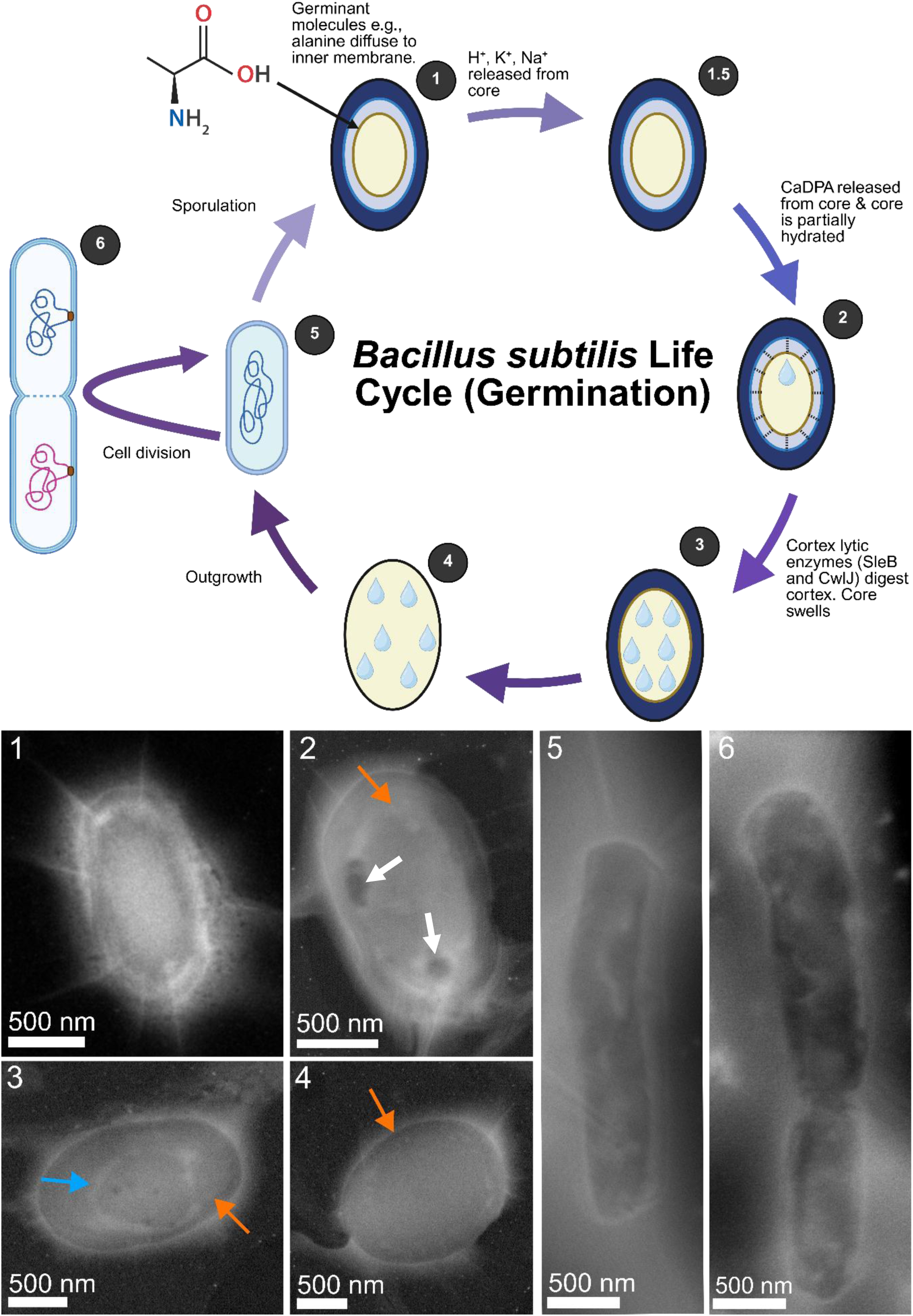
*B. subtilis* spore germination cycle showing internal structural changes. **(Top)** Schematic of the *B. subtilis* spore germination and outgrowth cycle, progressing clockwise from the dormant spore (1) to vegetative cell emergence (5), and cell division (6). Dormant spores (1) are metabolically inactive until exposure to a nutrient germinant such as L-alanine. The spore shown here is the same as in Figure 2E. Spores then begin germination (2), marked by partial core rehydration and cortex degradation (3). Continued rehydration leads to full germination (4) and outgrowth resulting in a vegetative *B. subtilis* cell (5), the normal life cycle then continues through bacterial cell division (6). **(Bottom)** Corresponding LPBSEM images showing representative internal features at each stage. (1) Dormant spore with bright, dense core and intact protective layers. (2–4) White arrows point to dark densities at the boundary of the core and cortex, the core membrane expands as germination progresses (orange arrow, 2-4) and intersects the cell wall in (4). In (2) there are two clear dark density regions due to a loss of material density. In (3), there is a bright ring of material (blue arrow) which is likely to be the nucleoid. (5 and 6) vegetative cells with internal detail visible such as DNA (5) and cell division (6).

Upon germination, clear morphological and contrast changes are observed. In an early stage, the overall morphology and shape of the spore remain largely unchanged, consistent with the initial stages of germination (Figure 4(2)). At this stage, remnants of the boundary between the core and cortex remain visible showing that core swelling is incomplete. Additionally, two localised regions of reduced contrast are observed around the core–cortex boundary (Figure 4(2), arrows). As BSE contrast is sensitive to material density, these features may reflect localised decreases in density. This observation is consistent with the possibility of early, spatially heterogeneous changes in the cortex during germination resulting from cortex degradation. The presence of one or two such regions is also consistent with previous reports of a limited number of germinosomes^71^ per spore, although no direct identification of these structures is possible in the present data. Importantly, core swelling can only occur if membrane vesicles present in the core are incorporated into the spore inner membrane^72^. Hence, if cortex hydrolysis precedes this step, it could create empty space within the spore that is then available to accommodate the expanding core.

At later stages of germination, spores progressively lose their distinct layered architecture, transitioning towards a more homogeneous internal structure. This change is consistent with continued cortex degradation and increasing core hydration. In Figure 4(3), the core appears to expand but has not yet reached the outer boundary (orange arrow), indicating that expansion is still ongoing. A central low-contrast structure is also observed (blue arrow), which may correspond to internal organisation within the core. While its position and morphology are broadly consistent with previously reported nucleoid localisation in germination^73^ this should been seen as tentative.

This demonstrates that LPBSEM provides a label-free method for visualising intracellular organisation during transitions between dormancy and germination. Finally, in the germination process, a fully germinated spore is observed (Figure 4(4)) with a swollen homogenous structure, and the core fully expanded to the outermost membrane.

Although we did not investigate outgrowth in this study, we have also imaged vegetative *B. subtilis* cells (Figure 4(5)) and vegetative cells during cell division (Figure 4(6)). In the vegetative cells, there is clear ultrastructural detail visible in the inner regions of the bacteria. This highlights that LPBSEM is not just useful for imaging spores but can also observe ultrastructural details from a range of samples without the need for potentially degrading sample preparation methods such as FIB milling, ultramicrotome slicing, heavy metal staining or cryogenic conditions.

Overall, these results demonstrate the value of LPBSEM for structural imaging and tracking of biological processes. By preserving hydration and minimising preparation artifacts, we have observed spore structural changes, as an initial platform for studying a wide range of dynamic events in microbiology at room temperature.

## Conclusion

We have used LPBSEM in an approach that combines graphene encapsulation with BSE-SEM to enable high-contrast imaging of hydrated biological specimens at room-temperature with minimal sample preparation. Specifically, we have imaged internal changes during spore germination, providing insights into potential cortex degradation and nucleoid organisation without the need for staining or fluorescent labels. Using *B. subtilis* spores as a model system, we show that this methodology preserves hydration, prevents artifacts such as shrinkage and collapse and reveals the internal structures of the spores including the core, cortex, and inner and outer coat. By varying the accelerating voltage, we have further shown that it is possible to modulate the imaging depth and contrast, confirmed by Monte Carlo simulations.

## Methods

### Sporulation of *B. subtilis*

Wildtype (WT) *Bacillus subtilis (B. subtilis)* strain PY79^74^ was obtained from the Christie lab (University of Cambridge). Vegetative *B. subtilis* cells were cultured in Luria Broth at 37°C with no antibiotics used. For spore preparation, vegetative *B. subtilis* cells were plated on 2x Schaeffer’s-glucose (SG) medium agar plates^75^ with no antibiotic and incubated at 37°C for five days. Spores were harvested and purified by repeated washing with ice cold water by centrifugation. The presence of spores was confirmed by phase contrast light microscopy (Supplementary Information: Figure S1).

### Germination of *B. subtilis* Spores

*B. subtilis* spores were initially heat-activated at 70°C for 30 min to induce break dormancy and improve germination competence. Following heat activation, spores were rapidly cooled on ice and resuspended in an L-alanine–containing germination buffer (10 mM L-alanine, 1 mM glucose, 1 mM NaCl, and 25 mM HEPES; pH 7.8). The spore suspension was incubated at 37 °C with shaking at 180RPM to initiate germination. Samples were graphene encapsulated at 5 minutes, 20 minutes, and 1 hour after the start of incubation for BSE-SEM imaging.

### Single-Crystalline Monolayer Graphene Fabrication

A polycrystalline Cu foil was thermally converted into a single-crystalline Cu(111) foil using a contact-free annealing method^76^. The Cu (111) foil was then loaded into a CVD chamber and heated to 1055 °C in a mixed flow of 300 sccm Ar and 40 sccm H₂ for 1 hour at a pressure of 30 Torr. After annealing, 35 sccm CH₄ (0.10% in Ar) was introduced for 1 hour to grow monolayer single-crystalline graphene. Following graphene growth, the hot chamber was removed to cool the Cu foil under the same gas mixture and pressure^77^.

### Graphene Encapsulation

Graphene TEM grids were prepared by transferring graphene to a TEM grid using cellulose acetate butyrate (CAB)^78^. The cellulose-based polymer was prepared by mixing CAB (Merck) with ethyl acetate. The polymer solution was then deposited on top of CVD graphene grown on copper (Graphenea Inc., Grolltex, and Ruoff Group) using a spin coater (Ossila Spin Coater) at 6000 rpm for 60s. Subsequently, the copper foil was floated on an aqueous solution of ammonium persulfate (3.0 g dissolved in 100 mL of ultrapure water) until the copper foil was completely etched. After etching, the solution was replaced with ultrapure water to remove the etching solvent over several cycles.

Holey carbon-coated gold TEM grids (Quantifoil, 200 mesh, holes 2 µm in diameter) were introduced into the water solution on a filter paper placed at the bottom of a liquid container. Water was carefully removed using a pipette until the CAB+graphene layer was deposited on top of the grids. Finally, the resulting graphene TEM grid was placed on activated carbon, heated to 310 °C, and left overnight to remove the polymer.

This process also allows the incorporation of additional layers by applying a gentle glow-discharge treatment (7 mA for 10 seconds) to render the previous graphene layers hydrophilic, enabling the next graphene layer to deposit on top.

The graphene TEM grids were subsequently cleaned of polymer, and the grids were placed at the bottom of a Petri dish with another bilayer graphene layer floating on a liquid solution, phosphate buffer (PB) buffer or water. The solution containing the dormant or germinated *B. subtilis* spores was introduced to the liquid environment between the bottom grids and the floating flakes. The liquid was then gently removed until the floating flake contacted the top of the graphene TEM grids enclosing the spores.

### SEM and Raman Spectroscopy

SEM imaging used a JIB-4700F Multi Beam System (JEOL, Japan) at the electron Physical Science Imaging Centre (ePSIC, Harwell) for room temperature SEM at either 5, 10 or 15 keV accelerating voltage. Both SE and BSE signals were collected simultaneously using a scan time of 20 seconds for 1280 x 960 pixels at a working distance of 6.0-8.5 mm. The recorded probe current was 250 pA. The secondary electron signal was collected using an Everhart-Thornley detector, while the backscattered electrons were collected by a retractable backscattered electron detector. The total fluences for each dataset are reported in the Supplementary Information: Table S1. Raman spectroscopy used a WITec GmbH instrument with a 488 nm laser at 1.0 mW (Supplementary Information: Figure S6).

### Image Processing

Brightness and contrast were adjusted in Digital Micrograph version 3.51.3720.0 (Gatan, AMETEK). No further adjustments or filters were applied to any images. Line profiles in Figure 2 were drawn as indicated on the figures and values of the counts against the length across the line profile were exported to excel and normalised as the lowest value in the profile subtracted from all other values. The line profiles were plotted using a python script that included a spline smoothing factor to reduce noise. A spline smoothing factor of 300 was chosen. Data shown in Figure 2 was processed using spline smoothing. Line profiles in Figure 3 were plotted directly in Excel.

### Monte Carlo Simulations

Monte Carlo simulations using CASINO V2.51^50^ were used to model electron interaction volumes at varying accelerating voltages and validate experimental imaging conditions and contrast.

The CASINO software allows the modelling of a sample based on thickness, composition and density. These are all parameters that contribute to electron backscattering. As the *B. subtilis* spore represents a complex biological structure, this modelling framework should be considered an informed approximation rather than an absolute representation, as natural variability exists between individual spores.

A total of 200,000 electrons were simulated with incident beam energies ranging from 1 to 15 keV. The sample was oriented at a tilt angle of 0°, and the electron beam radius was set to 1.2 nm for all simulations to replicate the experimental conditions.

## Acknowledgements

J.B. and A.P.-T. contributed equally to this work. The authors acknowledge that The Rosalind Franklin Institute is funded by the UK Research and Innovation and Engineering and Physical Sciences Research Council. Y.M. and R.S.R. are funded by the Institute for Basic Science (IBSR019-D1). This research did not receive any specific grant from funding agencies in the public, commercial, or not-for-profit sectors. Dr. Patricia Bondia (Scientific Visual Communication Studio) created the illustrations in Figure 1.

We thank Diamond Light Source for access and support in use of the electron Physical Science Imaging Centre (Instrument E03 and proposal number MG23286) that contributed to the results presented here.

We thank Dr. Marcus Gallagher-Jones for the input and valuable discussion of this work.

## References

1. Williams, D. B. & Carter, C. B. Transmission Electron Microscopy. (Springer US, Boston, MA, 2009).

2. Goldstein, J. I. et al. Scanning Electron Microscopy and X-Ray Microanalysis. (Springer, New York, NY, 2018).

3. Miranda, K., Girard-Dias, W., Attias, M., de Souza, W. & Ramos, I. Three dimensional reconstruction by electron microscopy in the life sciences: An introduction for cell and tissue biologists. Mol. Reprod. Dev. 82, 530–547 (2015).

4. Dykstra, M. J. & Reuss, L. E. Biological Electron Microscopy: Theory, Techniques, and Troubleshooting. (Springer US, Boston, MA, 2003).

5. Titze, B. & Genoud, C. Volume scanning electron microscopy for imaging biological ultrastructure. Biol. Cell 108, 307–323 (2016).

6. Kourkoutis, L. F., Plitzko, J. M. & Baumeister, W. Electron Microscopy of Biological Materials at the Nanometer Scale. Annu. Rev. Mater. Res. 42, 33–58 (2012).

7. Ayache, J., Beaunier, L., Boumendil, J., Ehret, G. & Laub, D. Sample Preparation Handbook for Transmission Electron Microscopy. (Springer, New York, NY, 2010).

8. Mollenhauer, H. H. Artifacts caused by dehydration and epoxy embedding in transmission electron microscopy. Microsc. Res. Tech. 26, 496–512 (1993).

9. McCaffrey, J. P., Phaneuf, M. W. & Madsen, L. D. Surface damage formation during ion-beam thinning of samples for transmission electron microscopy. Ultramicroscopy 87, 97–104 (2001).

10. Cazaux, J. Recent developments and new strategies in scanning electron microscopy. J. Microsc. 217, 16–35 (2005).

11. Ogura, K. & Hasegawa, Y. Application of Backscattered Electron Image to Biological Specimens. J. Electron Microsc. (Tokyo) 29, 68–71 (1980).

12. Koga, D., Bochimoto, H., Watanabe, T. & Ushiki, T. Backscattered electron image of osmium-impregnated/macerated tissues as a novel technique for identifying the cis-face of the Golgi apparatus by high-resolution scanning electron microscopy. J. Microsc. 263, 87–96 (2016).

13. Kariya, T. et al. Direct evidence for activated CD8+ T cell transmigration across portal vein endothelial cells in liver graft rejection. J. Gastroenterol. 51, 985–998 (2016).

14. Lucas, B. A. & Grigorieff, N. Quantification of gallium cryo-FIB milling damage in biological lamellae. Proc. Natl. Acad. Sci. U. S. A. 120, e2301852120 (2023).

15. Koga, D., Kusumi, S., Shibata, M. & Watanabe, T. Applications of Scanning Electron Microscopy Using Secondary and Backscattered Electron Signals in Neural Structure. Front. Neuroanat. 15, 759804 (2021).

16. Rice, W. J. et al. Routine determination of ice thickness for cryo-EM grids. J. Struct. Biol. 204, 38–44 (2018).

17. Thompson, R. F., Walker, M., Siebert, C. A., Muench, S. P. & Ranson, N. A. An introduction to sample preparation and imaging by cryo-electron microscopy for structural biology. Methods 100, 3–15 (2016).

18. Caffrey, B. J. et al. Liquid Phase Electron Microscopy of Bacterial Ultrastructure. Small 20, 2402871 (2024).

19. Pedrazo-Tardajos, A. et al. Direct visualization of ligands on gold nanoparticles in a liquid environment. Nat. Chem. 16, 1278–1285 (2024).

20. Arenas Esteban, D., et al. Quantitative 3D structural analysis of small colloidal assemblies under native conditions by liquid-cell fast electron tomography. Nat. Commun. 15, 6399 (2024).

21. Liu, K.-L. et al. Novel microchip for in situ TEM imaging of living organisms and bio-reactions in aqueous conditions. Lab. Chip 8, 1915–1921 (2008).

22. Yuk, J. M. et al. High-Resolution EM of Colloidal Nanocrystal Growth Using Graphene Liquid Cells. Science 336, 61–64 (2012).

23. Keskin, S. & de Jonge, N. Reduced Radiation Damage in Transmission Electron Microscopy of Proteins in Graphene Liquid Cells. Nano Lett. 18, 7435–7440 (2018).

24. Jonge, N. de, Peckys, D. B., Kremers, G. J. & Piston, D. W. Electron microscopy of whole cells in liquid with nanometer resolution. Proc. Natl. Acad. Sci. 106, 2159–2164 (2009).

25. Huang, T.-W. et al. Self-aligned wet-cell for hydrated microbiology observation in TEM. Lab. Chip 12, 340–347 (2012).

26. Kelly, D. J. et al. Nanometer Resolution Elemental Mapping in Graphene-Based TEM Liquid Cells. Nano Lett. 18, 1168–1174 (2018).

27. Algara-Siller, G., Kurasch, S., Sedighi, M., Lehtinen, O. & Kaiser, U. The pristine atomic structure of MoS2 monolayer protected from electron radiation damage by graphene. Appl. Phys. Lett. 103, 203107 (2013).

28. Cho, H. et al. The Use of Graphene and Its Derivatives for Liquid-Phase Transmission Electron Microscopy of Radiation-Sensitive Specimens. Nano Lett. 17, 414–420 (2017).

29. He, C. & Donald, A. M. Morphology of Core-Shell Polymer Latices during Drying. Langmuir 12, 6250–6256 (1996).

30. Zheng, T., Waldron, K. W. & Donald, A. M. Investigation of viability of plant tissue in the environmental scanning electron microscopy. Planta 230, 1105–1113 (2009).

31. Donald, A. M. The use of environmental scanning electron microscopy for imaging wet and insulating materials. Nat. Mater. 2, 511–516 (2003).

32. Kirk, S. E., Skepper, J. N. & Donald, A. M. Application of environmental scanning electron microscopy to determine biological surface structure. J. Microsc. 233, 205–224 (2009).

33. Driks, A. Overview: Development in bacteria: spore formation in Bacillus subtilis. Cell. Mol. Life Sci. 59, 389–391 (2002).

34. Zhang, X., Al-Dossary, A., Hussain, M., Setlow, P. & Li, J. Applications of Bacillus subtilis Spores in Biotechnology and Advanced Materials. Appl. Environ. Microbiol. 86, e01096–20 (2020).

35. Setlow, P. Spores of Bacillus subtilis: their resistance to and killing by radiation, heat and chemicals. J. Appl. Microbiol. 101, 514–525 (2006).

36. Setlow, P. Resistance of Bacterial Spores to Chemical Agents. in Russell, Hugo & Ayliffe’s 121–130 (John Wiley & Sons, Ltd, 2013).

37. Leggett, M. J., McDonnell, G., Denyer, S. P., Setlow, P. & Maillard, J.-Y. Bacterial spore structures and their protective role in biocide resistance. J. Appl. Microbiol. 113, 485–498 (2012).

38. Bauda, E. et al. Ultrastructure of macromolecular assemblies contributing to bacterial spore resistance revealed by in situ cryo-electron tomography. Nat. Commun. 15, 1376 (2024).

39. Goldman, R. C. & Tipper, D. J. Bacillus subtilis spore coats: complexity and purification of a unique polypeptide component. J. Bacteriol. 135, 1091–1106 (1978).

40. Laue, M., Niederwöhrmeier, B. & Bannert, N. Rapid diagnostic thin section electron microscopy of bacterial endospores. J. Microbiol. Methods 70, 45–54 (2007).

41. Waller, L. N., Fox, N., Fox, K. F., Fox, A. & Price, R. L. Ruthenium red staining for ultrastructural visualization of a glycoprotein layer surrounding the spore of Bacillus anthracis and Bacillus subtilis. J. Microbiol. Methods 58, 23–30 (2004).

42. Dierichs, R. Ruthenium red as a stain for electron microscopy. Some new aspects of its application and mode of action. Histochemistry 64, 171–187 (1979).

43. Blanquet, P. R. Ultrahistochemical study on the ruthenium red surface staining. Histochemistry 47, 63–78 (1976).

44. Isticato, R. & Ricca, E. Spore Surface Display. Microbiol. Spectr. 2, (2014).

45. Negri, A., Potocki, W., Iwanicki, A., Obuchowski, M. & Hinc, K. Expression and display of Clostridium difficile protein FliD on the surface of Bacillus subtilis spores. J. Med. Microbiol. 62, 1379–1385 (2013).

46. Xu, X. et al. Production of N-acetyl-D-neuraminic acid by use of an efficient spore surface display system. Appl. Environ. Microbiol. 77, 3197–3201 (2011).

47. Driks, A. & Eichenberger, P. The Spore Coat. Microbiol. Spectr. 4, (2016).

48. Freitas, C. et al. A protein phosphorylation module patterns the Bacillus subtilis spore outer coat. Mol. Microbiol. 114, 934–951 (2020).

49. Popham, D. L., Helin, J., Costello, C. E. & Setlow, P. Muramic lactam in peptidoglycan of Bacillus subtilis spores is required for spore outgrowth but not for spore dehydration or heat resistance. Proc. Natl. Acad. Sci. U. S. A. 93, 15405–15410 (1996).

50. Drouin, D. et al. CASINO V2.42—A Fast and Easy-to-use Modeling Tool for Scanning Electron Microscopy and Microanalysis Users. Scanning 29, 92–101 (2007).

51. Plomp, M., Leighton, T. J., Wheeler, K. E. & Malkin, A. J. The High-Resolution Architecture and Structural Dynamics of *Bacillus* Spores. Biophys. J. 88, 603–608 (2005).

52. Driks, A. Bacillus subtilis Spore Coat. Microbiol. Mol. Biol. Rev. 63, 1–20 (1999).

53. Abhyankar, W. R. et al. The Influence of Sporulation Conditions on the Spore Coat Protein Composition of Bacillus subtilis Spores. Front. Microbiol. 7, 1636 (2016).

54. Luo, J. et al. Ultralow Secondary Electron Emission of Graphene. ACS Nano 5, 1047–1055 (2011).

55. Velazco, A. et al. Reduction of SEM charging artefacts in native cryogenic biological samples. Nat. Commun. 16, 5204 (2025).

56. Zhang, Y. et al. Charging of Vitreous Samples in Cryogenic Electron Microscopy Mitigated by Graphene. ACS Nano 17, 15836–15846 (2023).

57. Kanaya, K. & Okayama, S. Penetration and energy-loss theory of electrons in solid targets. J. Phys. D: Appl. Phys. 5, 43 (1972).

58. Berlin, E., Curran, H. R. & Pallansch, M. J. Physical Surface Features and Chemical Density of Dry Bacterial Spores. J. Bacteriol. 86, 1030–1036 (1963).

59. Mao, L. et al. Protein Profile of Bacillus subtilis Spore. Curr. Microbiol. 63, 198–205 (2011).

60. Atrih, A. & Foster, S. J. Analysis of the role of bacterial endospore cortex structure in resistance properties and demonstration of its conservation amongst species. J. Appl. Microbiol. 91, 364–372 (2001).

61. Glauner, B. Separation and quantification of muropeptides with high-performance liquid chromatography. Anal. Biochem. 172, 451–464 (1988).

62. Kurniawan, O. & Ong, V. K. S. Investigation of range-energy relationships for low-energy electron beams in silicon and gallium nitride. Scanning 29, 280–286 (2007).

63. Moir, A. & Cooper, G. Spore Germination. Microbiol. Spectr. 3, TBS-0014–2012 (2015).

64. Christie, G. & Setlow, P. Bacillus spore germination: Knowns, unknowns and what we need to learn. Cell. Signal. 74, 109729 (2020).

65. Fiester, S. E., Helfinstine, S. L., Redfearn, J. C., Uribe, R. M. & Woolverton, C. J. Electron Beam Irradiation Dose Dependently Damages the Bacillus Spore Coat and Spore Membrane. Int. J. Microbiol. 2012, 579593 (2012).

66. Yi, X. & Setlow, P. Studies of the Commitment Step in the Germination of Spores of Bacillus Species. J. Bacteriol. 192, 3424–3433 (2010).

67. Hashimoto, T., Frieben, W. R. & Conti, S. F. Germination of Single Bacterial Spores. J. Bacteriol. 98, 1011–1020 (1969).

68. Setlow, B., Melly, E. & Setlow, P. Properties of Spores of Bacillus subtilis Blocked at an Intermediate Stage in Spore Germination. J. Bacteriol. 183, 4894–4899 (2001).

69. Gupta, P. et al. Early activation of bioenergetic metabolism powers bacterial spore germination. Proc. Natl. Acad. Sci. U. S. A. 122, e2510996122 (2025).

70. Paidhungat, M. & Setlow, P. Spore Germination and Outgrowth. in Bacillus subtilis and Its Closest Relatives 537–548 (John Wiley & Sons, Ltd, 2001).

71. Wen, J., Pasman, R., Manders, E. M. M., Setlow, P. & Brul, S. Visualization of Germinosomes and the Inner Membrane in Bacillus subtilis Spores. J. Vis. Exp. 146, e59388 (2019).

72. Laue, M., Han, H.-M., Dittmann, C. & Setlow, P. Intracellular membranes of bacterial endospores are reservoirs for spore core membrane expansion during spore germination. Sci. Rep. 8, 11388 (2018).

73. Ragkousi, K., Cowan, A. E., Ross, M. A. & Setlow, P. Analysis of Nucleoid Morphology during Germination and Outgrowth of Spores of Bacillus Species. J. Bacteriol. 182, 5556–5562 (2000).

74. Schroeder, J. W. & Simmons, L. A. Complete Genome Sequence of Bacillus subtilis Strain PY79. Genome Announc. 1, e01085–13 (2013).

75. Nicholson, W. & Setlow, P. Sporulation, germination and outgrowth. in Molecular Biological Methods for Bacillus 391–450 (John Wiley & Sons Ltd., 1990).

76. Jin, S. et al. Colossal grain growth yields single-crystal metal foils by contact-free annealing. Science 362, 1021–1025 (2018).

77. Luo, D. et al. Adlayer-Free Large-Area Single Crystal Graphene Grown on a Cu(111) Foil. Adv. Mater. 31, 1903615 (2019).

78. Jain, N. et al. Exploring the effects of graphene and temperature in reducing electron beam damage: A TEM and electron diffraction-based quantitative study on Lead Phthalocyanine (PbPc) crystals. Micron. 169, 103444 (2023).

