## Supplementary Information for "Liquid Phase Backscattered Scanning Electron Microscopy of *Bacillus subtilis* Spores"

**Electron beam damage**

The total electron fluence for each dataset was calculated using the equation:

$$Fluence= \frac{I \cdot t}{e^{-}\cdot A}\cdot N$$

Where *I* is the beam current, *t* is the dwell time, *e^-^* is the electron charge, *A* is the area of the pixel size and *N* is the number of frames. With a beam current of 250 pA, dwell time of 16.27 µs and pixel size as stated in table S1, the total fluences are presented below.

**Table S1**. Calculated electron fluence for each dataset.

| Dataset | Magnification (kX) | Pixel size  (nm) | Electron Fluence (e^-^/Å^2^) |
| --- | --- | --- | --- |
| Figure 2A, B | 17 | 5.9 | 7.3 |
| Figure 2D, E | 20 | 5.0 | 10.2 |
| Figure 2G, H | 17 | 5.9 | 7.3 |
| Figure 3A | 19 | 5.3 | 9.2 |
| Figure 3C | 23 | 4.4 | 13.4 |
| Figure 3E | 30 | 3.3 | 23.3 |
| Figure 4.1 | 20 | 5.0 | 10.2 |
| Figure 4.2, 4.3, 4.4 | 17 | 5.9 | 7.3 |
| Figure 4.5 | 22 | 5.5 | 8.6 |
| Figure 4.6 | 20 | 5.0 | 10.2 |
| Figure S1A | 16 | 6.3 | 6.5 |
| Figure S1B, C | 23 | 4.4 | 13.4 |
| Figure S2A-D | 27 | 3.7 | 13.7 |
| Figure S2E, F | 22 | 5.5 | 8.6 |
| Figure S2G, H | 23 | 4.4 | 13.4 |
| Figure S3A-D | 17 | 5.9 | 7.3 |
| Figure S4A, B | 8 | 12.5 | 1.6 |

When imaging biological specimens in low-voltage SEM as is done here, biological structures will undergo damage mechanisms such as radiolysis even at fluences of single digit e^-^/Å^2^ as only a few eV is needed to break molecular bonds^1,2^. In low-voltage image acquisition, the interaction volume is decreased meaning that damage may be localised to the top surface of the sample. Therefore, even modest exposures of a few electrons/Å² must be treated as part of the total “dose budget” to preserve fine structure in biological structures.

In our case, damage can be visualised by occasional bubbling of the liquid around and above the sample (Figure S1A) and shrinkage of the sample diameters (Figure S1B-C).

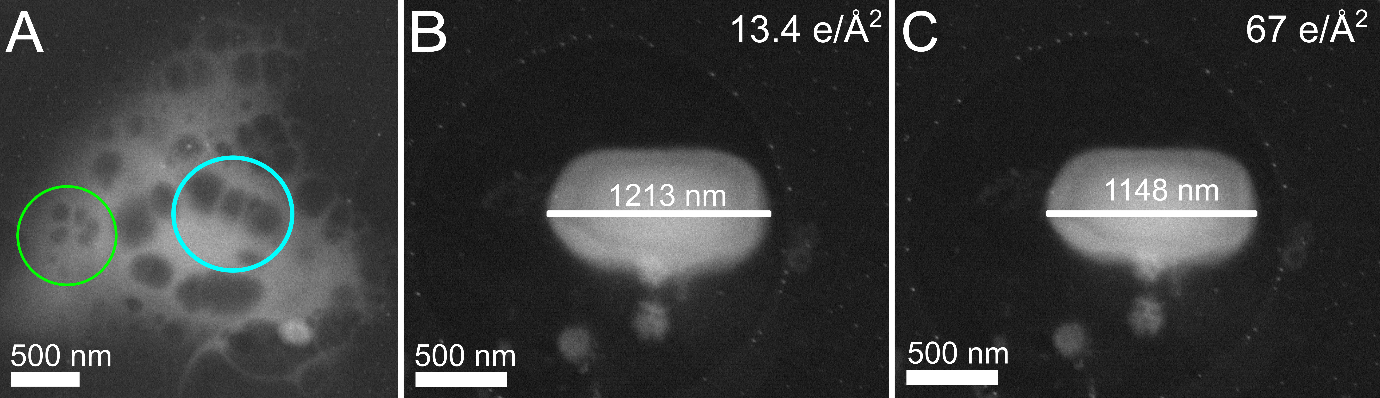
**Figure S1: Electron Beam Damage.** Electron beam damage can mainly manifest in 2 distinct ways. 1) Bubbling of the liquid as water bonds break and H_2_ and O_2_ gases are formed as shown in (A) and 2) the sample shrinking in size (B-C). Bubbling generally occurs when there is a relatively large volume of liquid, in this case PB buffer, in the GLC. The green circle highlights the bubbles that form in the GLC away from the sample whereas the light blue circle shows the bubbles forming above the sample. Panel B is an image of a *B. subtilis* spore imaged for the first time (13.4 e^-^/Å^2^). Panel C is an image of a *B. subtilis* spore that has been imaged 5 times (total fluence of 67 e^-^/Å^2^).

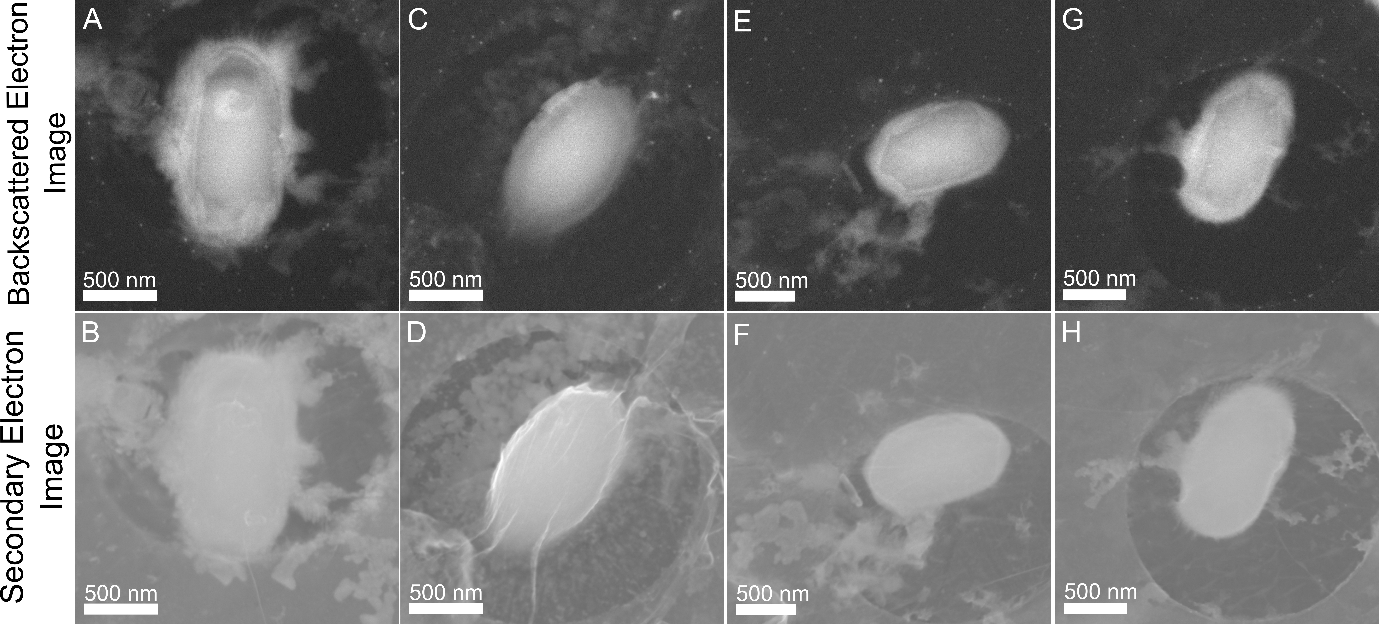

**Figure S2. *Bacillus subtilis* spores encapsulated in pure water and imaged using scanning electron microscopy (SEM).** The top row shows backscattered electron (BSE) images, while the bottom row shows secondary electron (SE) images, both acquired at an accelerating voltage of 10 keV.

**Graphene Backscatter Yield**

The backscattering coefficient (η), that is the ratio of backscattered electrons (BSE) (>50 eV) to incident electrons, is traditionally considered to scale with atomic number (Z), producing higher signal intensities in regions of higher-Z or higher density materials. However, this relationship becomes non-linear at lower beam energies (<5 keV), where η may increase for low-Z materials and decrease for high-Z ones^3,4^, limiting the applicability of conventional models such as the Everhart approximation. These deviations highlight the need for more accurate descriptions of BSE behaviour in low-voltage, surface-sensitive imaging regimes.

The backscatter yield of graphene was simulated using Monte Carlo simulations from 1 to 15 keV to model whether the graphene contributes to BSE images (Table S1). The graphene specimen was synthesised on CASINO simply by inputting a pure carbon sample of 0.35 nm in thickness and using the same microscope parameters as outline in the methods section.

**Table S2: Backscatter coefficient of a single layer of graphene at various accelerating voltages.**

| Accelerating Voltage (keV) | Backscatter coefficient (η) |
| --- | --- |
| 1 | 2.8x10^-3^ |
| 2 | 7.1x10^-4^ |
| 3 | 2.9x10^-4^ |
| 4 | 1.7x10^-4^ |
| 5 | 1.2x10^-4^ |
| 6 | 4.0x10^-5^ |
| 7 | 2.7x10^-5^ |
| 8 | 7.0x10^-6^ |
| 9 | 3.3x10^-5^ |
| 10 | 7.0x10^-6^ |
| 11 | 1.3x10^-5^ |
| 12 | 1.3x10^-5^ |
| 13 | 1.3x10^-5^ |
| 14 | 7.0x10^-6^ |
| 15 | 1.3x10^-5^ |

The backscatter yield using accelerating voltages of 1 – 15 keV, equivalent to those used in our experimental procedures are minimal highlighting that graphene will likely not contribute significantly to the final image. This also correlates with results in Figure 3G that shows the Monte Carlo simulations of the backscattered electron generation with the sample. In this figure, we see that the majority of BSEs are generated deeper in the sample.

**Incomplete Graphene Cover**

In the graphene encapsulation preparation, often we prepare a setup of a TEM grid, followed by two layers of graphene, sample, and then two additional layers of graphene above to “seal” the sample. We refer to this set up as the “double layer” approach where there are four sheets of graphene in total i.e., one set of two layers below the sample and two above. The reason for this being that if one layer of the graphene is to break during, for instance, grid handling, there is a greater chance the second layer will stay intact, keeping the sample encapsulated.

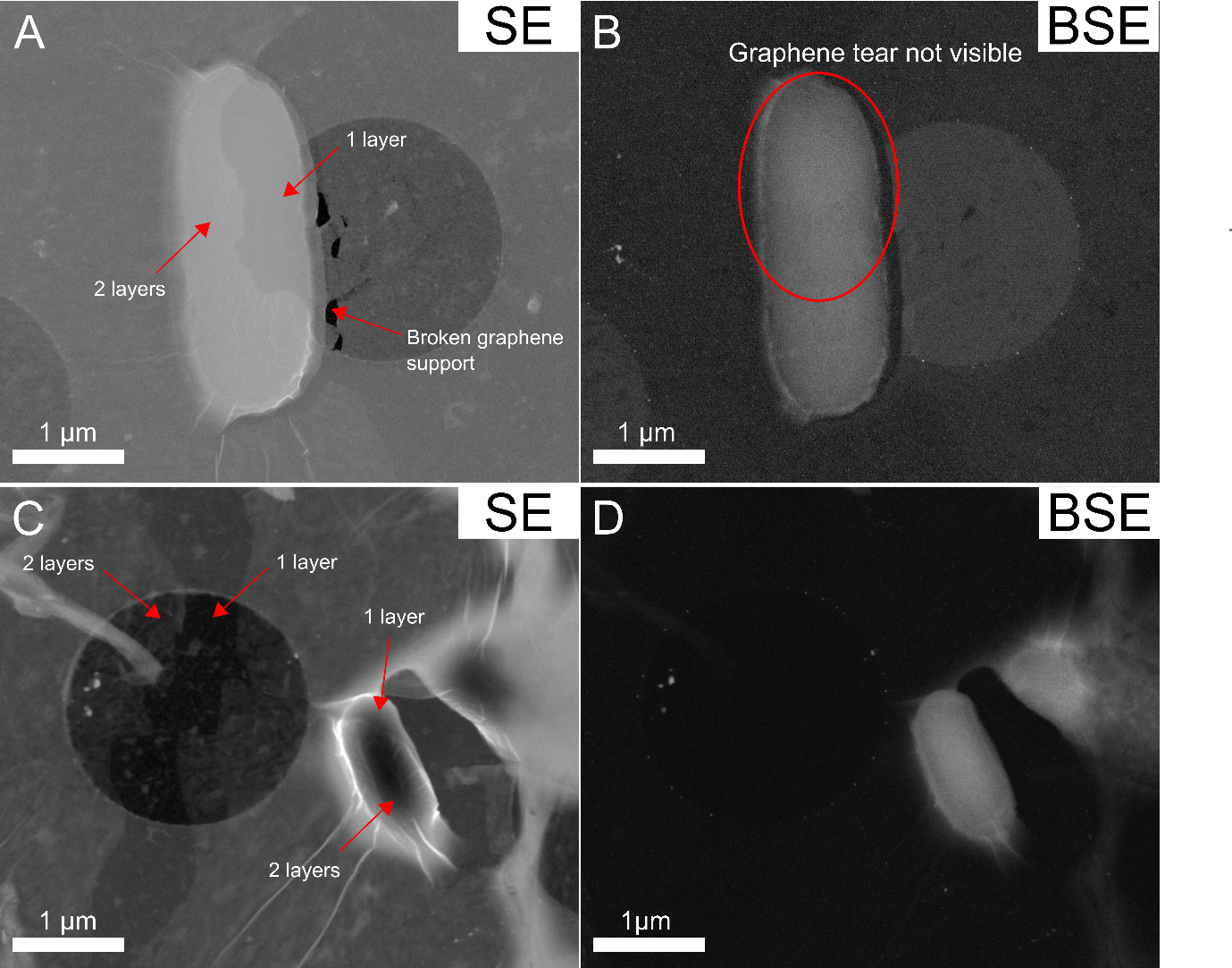

**Figure S3: SE and BSE SEM images of *E. coli* (A and B, respectively) and a *B. subtilis* spore (C and D, respectively) with incomplete encapsulation.** Areas of incomplete encapsulation can clearly be seen in the SE images with both samples, yet this is much less clear in the BSE. Panels A and B were imaged at 10 keV whereas C and D were imaged at 5 keV.

In Figure S3, the SE reveals for both samples that that the double layer graphene encapsulation is incomplete with visible cracks seen on and around the *E. coli* and *B. subtilis* spore. Usually, these regions would be avoided for imaging as local dehydration could occur. However, this highlights the use of both using double layer graphene and collecting the SE signal. Firstly, by using double layer graphene, if there is a break in the top layer, the second layer is there to provide further protection and prevent dehydration. Second, without collecting the SE signal, it would not be clear that the graphene was broken. First instance in Figure S3 panels A and B, the graphene tear on the *E. coli* cell is clear. This makes sense as we know that the SE signal gives us surface level information. However, since the bulk of the BSE signal comes from beneath the surface, the surface level defect is not visible in Figure S3B; this further supports the idea that the graphene contributes minimally to the BSE signal. If there was a significant contribution by graphene to the BSE, the break in the top layer would be clearly visible. Despite there being inconsistencies in the graphene, both the *E. coli* cell and *B. subtilis* spore have clearly defined membranes.

**Low magnification images/wide field of view**

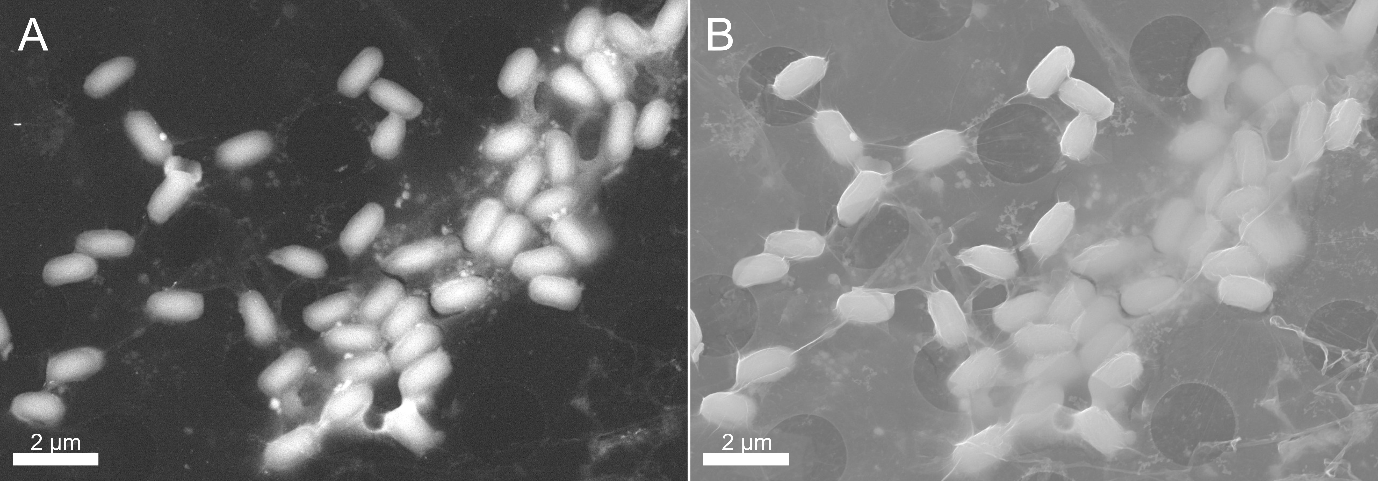

**Figure S4: Wide field of view BSE (a) and SE (b) images of graphene encapsulated *B. subtilis* spores.** The low-magnification image showing a representative region of the grid with the SE highlighting graphene coverage and BSE showing sample preservation. Spores were encapsulated in pure water and imaged at 10 keV.

To provide context for the higher-resolution images presented in the main text, Figure S3 shows representative wide-field views of the graphene-encapsulated *B. subtilis* spores in liquid. The low-magnification BSE images reveal that the graphene coverage can remain continuous over tens of micrometres, enabling imaging over a large field of view without significant charging or membrane rupture. However, this is not always the case, and we recommend that low magnification images should be take prior to high magnification images at various region to assess the graphene coverage across the grids and identify better and worse regions. Variations in the apparent backscattered intensity across the field may arise primarily from local thickness fluctuations in the liquid in the GLC. The wide field of view approach would be ideal in the future for large samples, e.g., imaging cell-cell interactions, and integrating in CLEM approaches. The ability to image in wide field of view in the SEM provides additional benefits to TEM as a sample can be efficiently screened for quality in an SEM with a relatively low fluence, particularly with the wide field of view approach. The calculated fluence in Figure S5 was just 1.6 e^-^/Å^2^.

**Phase Contrast Light Microscopy**

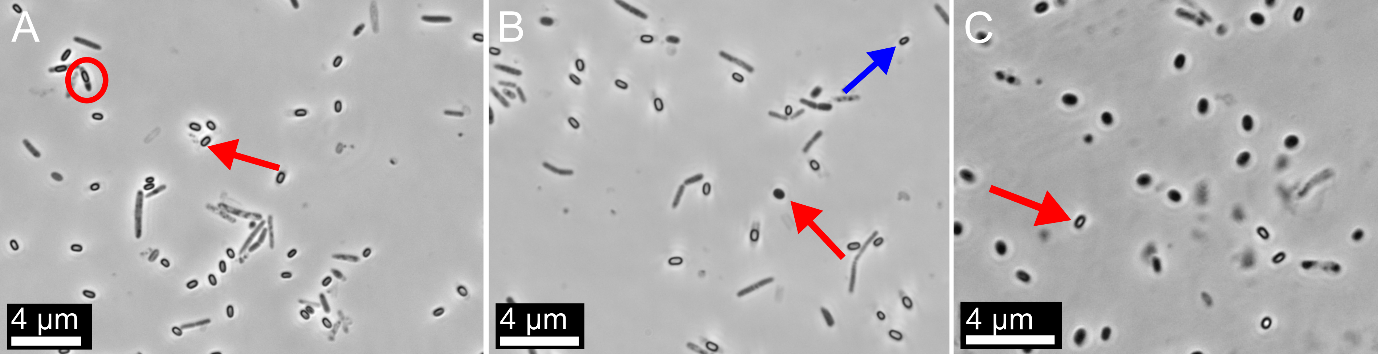

**Figure S5: Overview of changes observable by phase contrast light microscopy.** Prior to the initiation of germination, or immediately after, the population will look as in panel A. Spores have an elongated shape of ~1 µm in length with a phase bright centre (red arrow). Circled in red is a spore within a mother cell; it is worth noting that often spore populations are heterogenous in life cycle stage. In panel B, very soon post germination, the overall population is largely still mature spores, but some spores present with a less bright centre region (blue arrow). Additionally, some spores appear to have completed the germination process as they completely lose the phase bright centre (Panel B, red arrow) due to the loss of Calcium dipicolinic acid (CaDPA). Note here in panel B that the surrounding spores still have their phase bright centre indicating a heterogenous population with spores likely at different stages of germination. Therefore, it appears that germination in an entire population is difficult to control and induce at once. Finally, the vast majority of spores become phase dark indicating an end point for germination (Panel C).

**Graphene Characterisation**

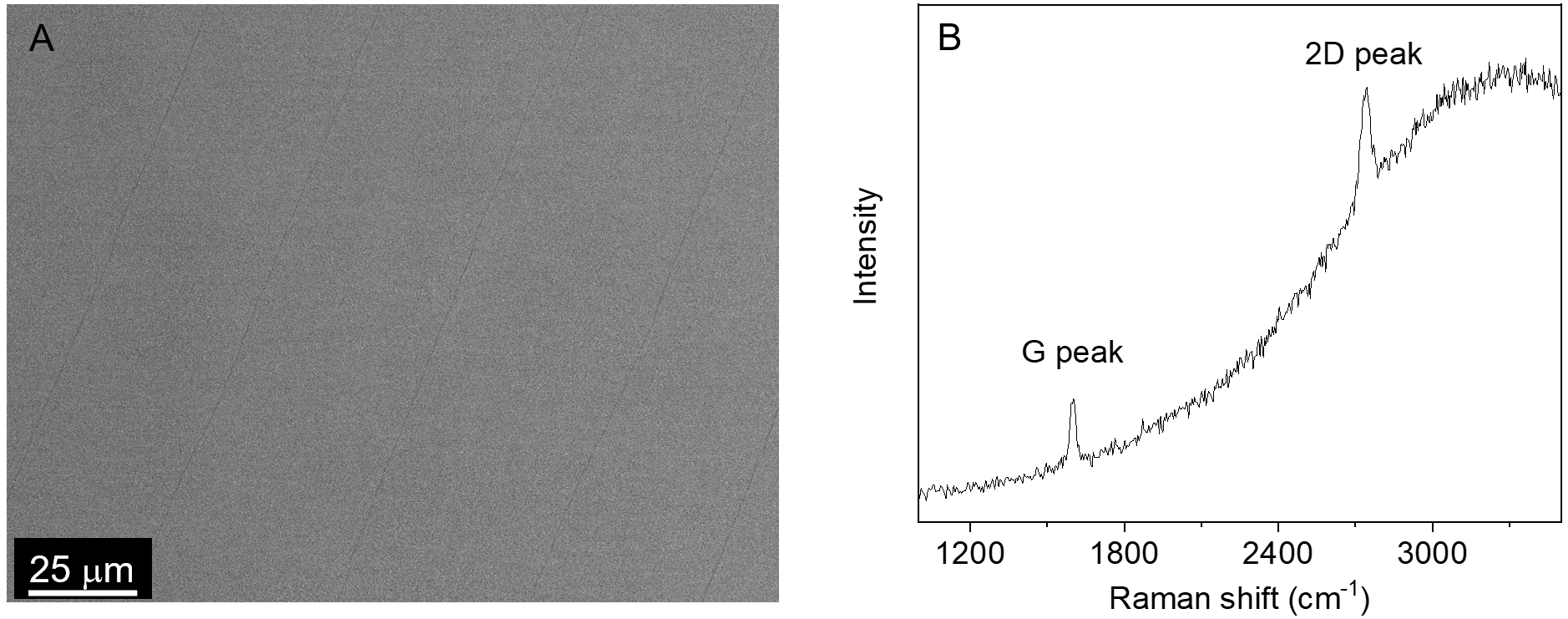

**Figure S6:** Evaluation of the quality of the single-crystal monolayer graphene. A, SEM image of the graphene film grown on a Cu(111) foil, showing a highly uniform surface morphology over a large area. B, Raman spectrum of the graphene on Cu(111), exhibiting a typical monolayer signature with a 2D to G intensity ratio of approximately 2 and no detectable D peak, indicating high structural quality.

**References**

1.	Pawley, J. B. LVSEM for Biology. in *Biological Low-Voltage Scanning Electron Microscopy* (eds Schatten, H. & Pawley, J. B.) 27–106 (Springer, New York, NY, 2008).

2.	Glaeser, R. M. Limitations to significant information in biological electron microscopy as a result of radiation damage. *J. Ultrastruct. Res.* **36**, 466–482 (1971).

3.	Čalkovský, M., Müller, E. & Gerthsen, D. Quantitative analysis of backscattered-electron contrast in scanning electron microscopy. *J. Microsc.* **289**, 32–47 (2023).

4.	Goldstein, J. I. *et al.* Backscattered Electrons. in *Scanning Electron Microscopy and X-Ray Microanalysis* (eds Goldstein, J. I. et al.) 15–28 (Springer, New York, NY, 2018).
